# ClonArch: Visualizing the Spatial Clonal Architecture of Tumors

**DOI:** 10.1101/2020.04.06.027912

**Authors:** Jiaqi Wu, Mohammed El-Kebir

## Abstract

**Motivation:** Cancer is caused by the accumulation of somatic mutations that lead to the formation of distinct populations of cells, called clones. The resulting clonal architecture is the main cause of relapse and resistance to treatment. With decreasing costs in DNA sequencing technology, rich cancer genomics datasets with many spatial sequencing samples are becoming increasingly available, enabling the inference of high-resolution tumor clones and prevalences across different spatial coordinates. While temporal and phylogenetic aspects of tumor evolution, such as clonal evolution over time and clonal response to treatment, are commonly visualized in various clonal evolution diagrams, visual analytics methods that reveal the spatial clonal architecture are missing.

**Results:** This paper introduces ClonArch, a web-based tool to interactively visualize the phylogenetic tree and spatial distribution of clones in a single tumor mass. ClonArch uses the marching squares algorithm to draw closed boundaries representing the presence of clones in a real or simulated tumor. ClonArch enables researchers to examine the spatial clonal architecture of a subset of relevant mutations at different prevalence thresholds and across multiple phylogenetic trees. In addition to simulated tumors with varying number of biopsies, we demonstrate the use of ClonArch on a hepatocellular carcinoma tumor with ~280 sequencing biopsies. ClonArch provides an automated way to interactively examine the spatial clonal architecture of a tumor, facilitating clinical and biological interpretations of the spatial aspects of intratumor heterogeneity.

**Availability:** https://github.com/elkebir-group/ClonArch

## 1 Introduction

Repeated and unchecked somatic mutations in cancer destabilize cells and lead to tumorigenesis, progression, and ultimately metastasis (Nowell, 1976). During tumorigenesis distinct cell populations, or *clones*, that accumulate a distinct set of mutations, arise from an evolutionary process (Fig. 1a). This phenomenon of *intra-tumor heterogeneity* is the main cause of relapse and resistance to treatment (Fisher *et al*., 2013; Tabassum and Polyak, 2015). Effective visual exploration by experts is crucial for the extraction of relevant information from cancer genomics data, including the discovery of rare genomic events, verification of data quality, or identification of key players in cancer development (Schroeder *et al*., 2013).

**Fig. 1:**
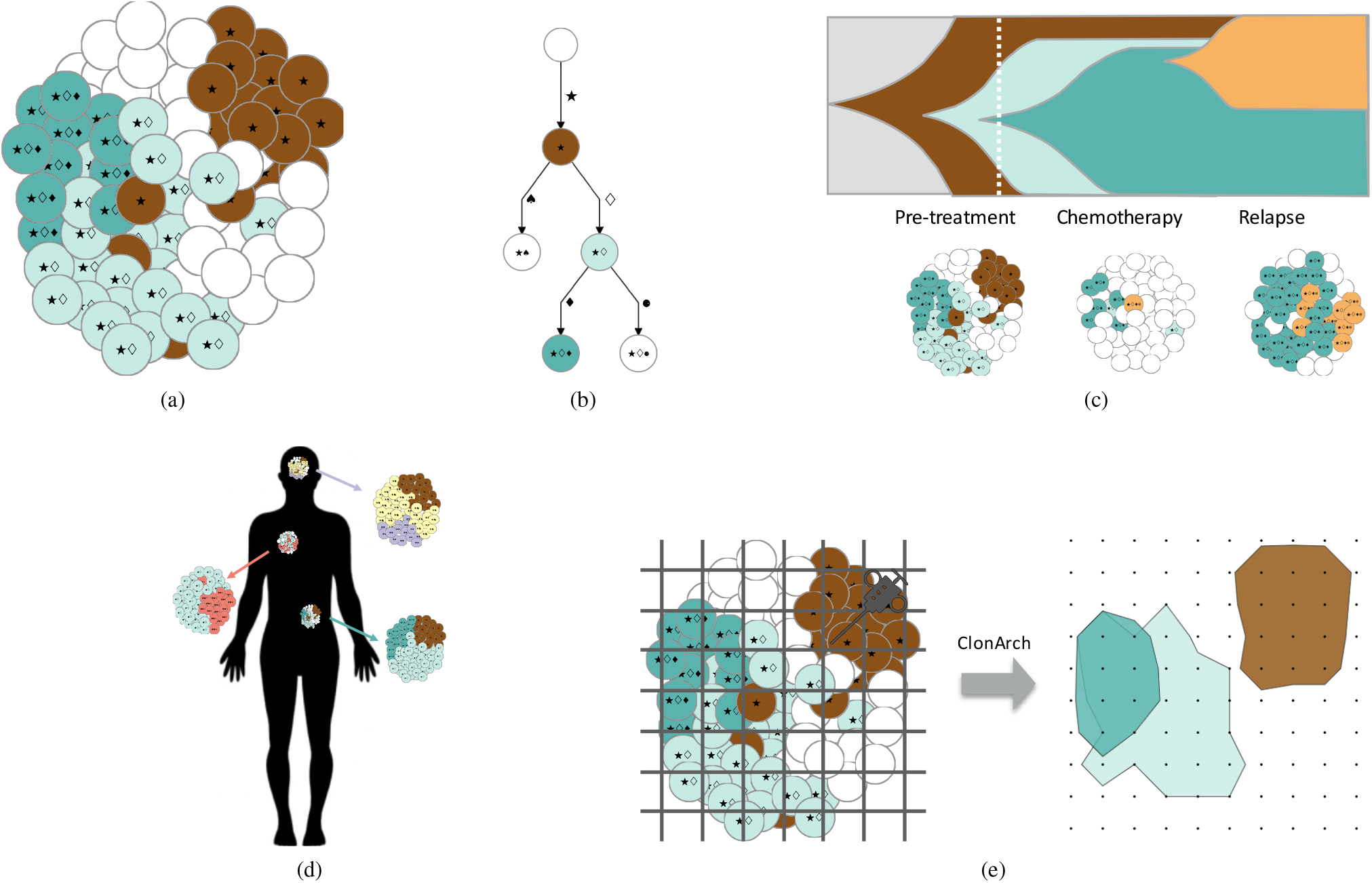
Overview of current and proposed tools for visualizing the clonal architecture of tumors. (a) A tumor is made up of multiple clones, each of which contains a distinct set of mutations. (b) Phylogenetic trees portray the evolutionary relationships between clones in a tumor. (c) Temporal representations of cancer evolution show tumor growth and clinical response. In addition, methods such as fishplot Miller *et al*., 2016 and TimeScape Smith *et al*., 2017 capture the evolutionary relationships between clones. (d) Presently, spatial representations of cancer evolution, obtained by methods such as MapScape Smith *et al*., 2017, illustrate clonal prevalences in different parts of the body after metastasis but do not show exact locations of clones within each anatomical location. (e) Given a phylogenetic tree and clonal prevalences at each sequencing biopsy, ClonArch represents the spatial distribution of clones on a grid using filled boundaries.

Recently, there has been a rise of visual analytics tools developed to analyze intra-tumor heterogeneity inferred from bulk or single-cell DNA sequencing data of tumors. Visualizations showing changes in clonal structure over time have been widely explored (Smith *et al*., 2017; Miller *et al*., 2016; Krzywinski, 2016). The most basic visualization is a phylogenetic tree, whose leaves correspond to tumor cells at the present time (clones), and whose edges are labeled by somatic mutations (Fig. 1b). While a phylogenetic tree provides a qualitative view of tumor progression, it does not show the prevalence or abundance of each clone. To overcome this limitation, the fishplot package for R enables the creation of temporal diagrams, previously done manually in vector-art programs, in an automated fashion (Miller *et al*., 2016). Specifically, fishplot estimates subclonal prevalence at multiple time points (e.g. pretreatment, post-treatment and relapse), and outputs a chart that represents clones at their relative proportions (Fig. 1c). Similarly, Smith *et al*. (2017) developed TimeScape, an automated tool that plots clonal prevalences (vertically) across time points (horizontally) for each clone. Diagrams of this type support illustration of details from broad trends in evolution or population dynamics of a few clones (Kvitek and Sherlock, 2013). However, while capturing the temporal and phylogenetic aspects of the© Wu and El-Kebir, 2020.tumor, these diagrams do not show spatial characteristics. MapScape, which comes from the same suite as TimeScape, is able to visualize spatially distinct tumor samples and indicate them on an anatomical image, or body map (Smith *et al*., 2017). MapScape uses colors to map clones to a phylogenetic tree that depicts their evolutionary relationships. Additionally, clonal composition per anatomical site is proportional to the corresponding clone’s colored region in the spatial representation (Fig. 1d). Although MapScape portrays a form of spatial visualization by representing clonal composition according to its prevalence, it does not convey the precise locations of each clone in an individual tumor.

Cancer genomics datasets with many spatial/regional sequencing biopsies from the same tumor are becoming increasingly available (Alves et *al*., 2019; Ding *et al*., 2019; Mamlouk *et al*., 2017; Ling et *al*., 2015; Gerlinger *et al*., 2012, 2014). Using multi-sample cancer phylogeny inference methods (El-Kebir *et al*., 2015; Malikic *et al*., 2015; Deshwar *et al*., 2015; Popic *et al*., 2015), such datasets enable researchers to infer detailed information on the spatial clonal architecture of individual tumors. Yet there is currently no method to visualize the spatial structure of a tumor that is, the distribution of clones at different spatial coordinates within a tumor. Analyzing the clonal composition of a tumor across space may facilitate clinical and biological interpretations and understanding of treatment resistance and cancer progression. For instance, structural information can be important for estimating the amount of genetic and clonal diversity in the tumor, and identifying the spatial relationships between driver mutations (Schroeder *et al*., 2013). Additionally, the introduction of spatially explicit population genetic models, which attempt to explain the variety of patterns observed in tumor architecture (Noble *et al*., 2019), calls for spatial visualization tools that can validate different models of tumor evolution.

Here, we fill the gap in available spatial composition tools by visualizing the spatial distribution of clones, by location, for a single tumor mass. We introduce ClonArch, a web-based method to interactively visualize tumor spatial structure given a set of phylogenetic trees and clone prevalences at distinct biopsies. From sequencing biopsy samples, we use cancer phylogeny inference methods (El-Kebir *et al*., 2015; Malikic *et al*., 2015; Deshwar *et al*., 2015; Popic *et al*., 2015) to obtain phylogenetic trees. Our proposed visual analytics approach uses the marching squares algorithm (Lorensen *et al*., 1987) to draw enclosed boundaries around clones above a specified threshold at each spatial location (Fig. 1e). We use ClonArch to analyze the spatial composition of a published human hepatocellular carcinoma composed of ~280 biopsies (Ling *et al*., 2015). In addition, we assess the applicability of ClonArch to datasets with fewer biopsies using simulations. ClonArch enables researchers to study the spatial aspects of intra-tumor heterogeneity, facilitating clinical and biological interpretations.

## 2 Requirements

Our aim is to develop a visual analytics tool to represent the spatial composition of a tumor in terms of clones and their prevalences. To obtain such a tool, we need to characterize our input data (Section 2.1) as well as typical visual analytics tasks (Section 2.2).

### 2.1 Data Characteristics

Our input data have the following characteristics:

### (D1) Mutations are grouped into clusters

Cancer phylogenetics pipelines group mutations that co-occur and never appear separate from each other into clusters that is, if all clones in the dataset that contain mutation *A* also contain mutation *B*, the two mutations are clustered together. Specialized methods exist for inferring mutation clusters from singlecell (Roth *et al*., 2016) and bulk (Roth *et al*., 2014; Miller *et al*., 2014) DNA sequencing data.

### (D2) There are multiple mutation clusters per clone

Clones are characterized by the accumulation of distinct mutations; a tumor will amass multiple mutation clusters as it evolves.

### (D3) Clones are distinguishable by mutation clusters

Each clone has a unique set of mutation clusters and thus mutations.

### (D4) Non-gridlike pattern of biopsy locations

To infer the spatial composition and evolutionary history of a tumor, we take multiple biopsies of a tumor, recording the 2-D spatial coordinate (*x, y*) of each location. Biopsy locations may not adhere to a perfect grid-like pattern.

### (D5) Clonal evolution described by a phylogenetic tree

From sequencing data of biopsies, we use cancer phylogeny methods specialized for singlecell (Ross and Markowetz, 2016; Jahn *et al*., 2016; El-Kebir, 2018) or bulk (El-Kebir *et al*., 2015; Malikic *et al*., 2015; Deshwar *et al*., 2015; Popic *et al*., 2015) DNA sequencing data to infer a set 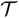 of phylogenetic trees *T*. The vertices of *T* correspond to tumor clones, and edges of *T* are labeled by mutation clusters.

### (D6) Clonal composition known at every biopsy

For each identified tree *T* in 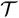, cancer phylogeny methods derive the proportion, or prevalence, of each clone in each sequencing biopsy. Due to intra-tumor heterogeneity, a single biopsy typically contains multiple clones.

## 2.2 Analysis Tasks

We identify the following analysis tasks.

### (A1) Spatial distribution of tumor clones

The user should be able to inspect the spatial distribution of tumor clones.

### (A2) Phylogenetic relationships of tumor clones

The clones of a tumor arise from an evolutionary process. The user should be able to inspect the phylogenetic tree(s) that relates the tumor clones.

### (A3) Ambiguity of tumor clones described by multiple phylogenetic trees

Typically, more than one phylogenetic tree can be inferred from the same sequencing data (Jamal-Hanjani *et al*., 2017; Pradhan and El-Kebir, 2018). If the data can be described by multiple trees, the user should be able to observe the spatial consistency between visualizations from different trees.

### (A4) Spatial distribution and phylogenetic relationships of clones restricted to a subset of relevant mutations

The majority of somatic mutations are passenger mutations that do not confer a selective advantage to the tumor as opposed to driver mutations. Similarly, not every somatic mutation is a drug target. Thus, for clinical and/or biological reasons, the user should be able to restrict the analysis to a subset *Z* of relevant somatic mutations. Original tumor clones that are identical with respect to this subset *Z* should be indistinguishable in the visualization.

### (A5) Relationship between prevalence and spatial distribution of an individual tumor clone

The user should be able to visually study the relationship between prevalence of a single tumor clone and spatial location. That is, the user should be able to identify the spatial locations in which a clone of interest occurs at a given prevalence.

### (A6) The relationship between prevalence and spatial distribution of multiple tumor clones

Analysis task (A5) should be extendable to multiple clones. That is, the user should be able to identify the spatial locations in which multiple clones of interest occur at possibly distinct prevalences.

### (A7) Exporting vector graphics of visualizations

The user should be able to export high-quality vector graphics of both the phylogenetic tree and spatial distribution of selected clones/mutations.

## 3 Methods

Section 3.1 formally defines the input to ClonArch according to the data characteristics specified in the previous section. Section 3.2 covers the analysis tasks pertaining to the phylogenetic tree, and the spatial distribution analysis tasks are described in Section 3.3. Finally, Section 3.4 introduces our visual analytics method that adheres to the outlined data characteristics and supports the identified analysis tasks.

### 3.1 Input

Our input is composed of a set 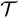 of phylogenetic trees (D5) and a set [*m*] = {1,…,*m*} of spatial biopsies (D6). A phylogenetic tree T is a tree with *n* = |*V*(*T*) | vertices rooted at vertex *r*(*T*). Each vertex *v* ∈ *V*(*T*) corresponds to a clone. Each edge (*v,w*) ∈ *E*(*T*) is labeled by one or more mutation clusters (D1), indicating the mutations that distinguish clone v from clone w (D2). Each clone *v* ∈ *V*(*T*) is composed of exactly those mutations that occur in the clusters that label the edges of the unique path from *r*(*T*) to *v* (D3). As such, the root vertex *r*(*T*) corresponds to the normal clone, not containing any mutations.

We know the spatial coordinates *σ*(*p*) = (*x,y*) of each biopsy *p*. Note that the set of spatial coordinates of all biopsies may not form a perfect grid (D4). Moreover, we know the clonal composition of every biopsy *p*. That is, we are given an *m* × *n* prevalence matrix *U* = [*u_pv_*],whereeach entry *u_p,v_* describes the prevalence of clone *v* in biopsy *p*. More formally, for each biopsy *p*, we have Σ_*v*∈*v*(*T*)_ *u_pv_* = 1 and *u_p,v_* ≥ 0 for each clone *v* (D6).

### 3.2 Phylogenetic Tree

Visualization. To accommodate analysis task A2 and showcase the phylogenetic relationship between clones, we use `dagre-d3`^1^ to draw a phylogenetic tree (Fig. 2). The root of the tree is the normal clone with no mutations, and the first edge contains the founding/trunk mutation(s). Each mutation cluster is assigned a symbol. Each vertex represents a clone, characterized by a color and a set of symbols representing the mutation clusters that make up the clone (D2). Each edge represents a newly introduced mutation cluster, and is labeled by the mutation names and corresponding symbol (D3). We limit the number of displayed labels per edge to six labels.

**Fig. 2:**
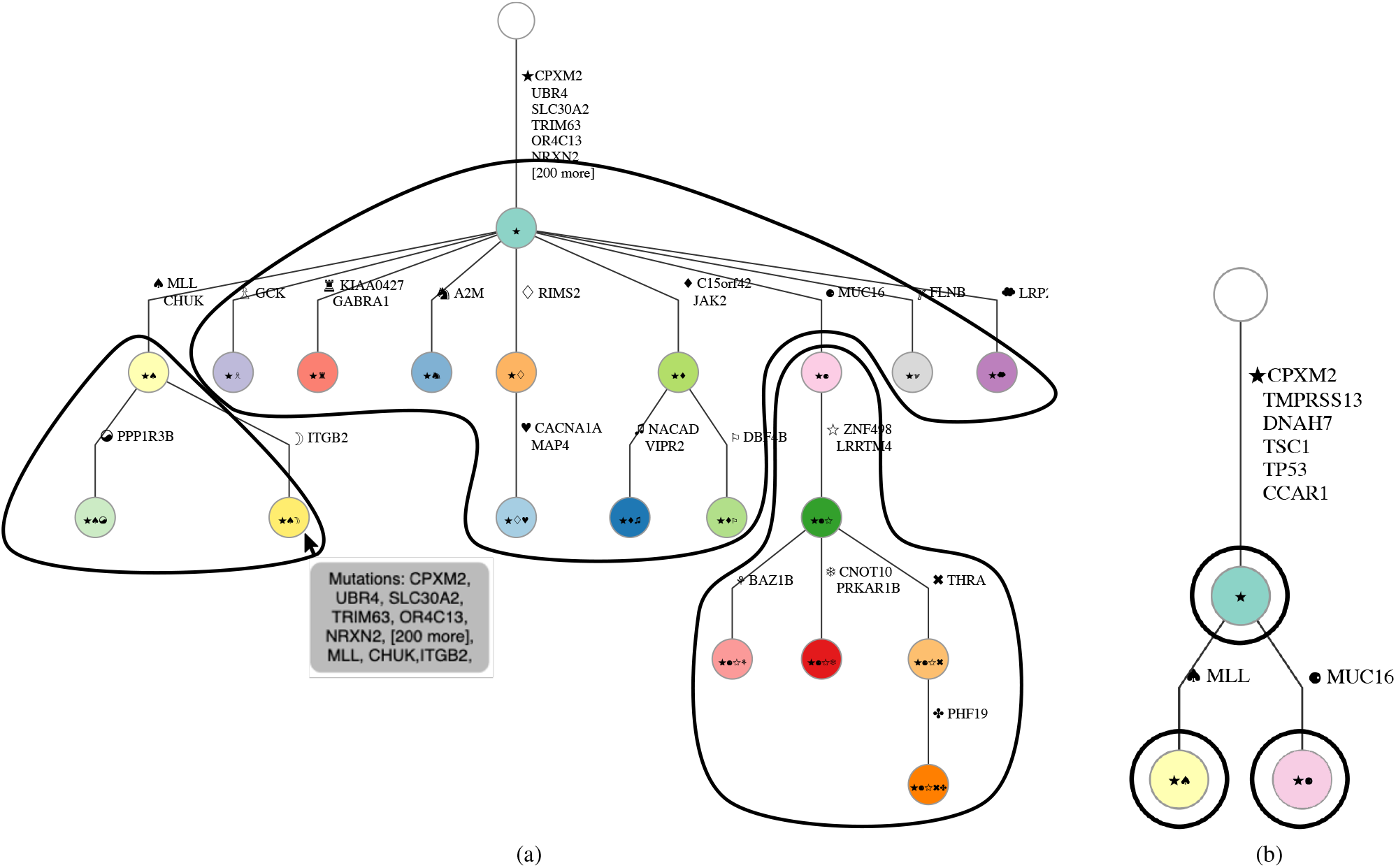
Phylogenetic tree describing relationship between clones in ClonArch. (a) Vertices are clones, distinguished by mutation clusters (symbols) that are introduced on the edges. In addition to displaying the mutation cluster symbol, each edge displays up to 6 mutations present in the cluster. Hovering over a vertex displays a tool tip listing the mutations in the corresponding clone. (b) Restricting our analysis to a subset of eight mutations (shown on the edge labels), results in a smaller phylogenetic tree, whose clones encompass multiple original clones.*Interactivity*. A drop down menu allows the user to select a tree to visualize, if there are multiple trees that describe the tumor data. Once selected, on hover, tree vertices will be drawn with a thicker stroke width and display a tool tip that summarizes the distinct mutations in that clone (Fig. 2). In addition, the corresponding clone on the grid is brought to the foreground. Clicking on a vertex allows a user to show and hide the corresponding clone on the grid. The fill color of a vertex is set to white when hiding the clone; the vertex regains the original color upon re-enabling the clone. Hovering over the vertex highlights the corresponding clone on the grid.

### 3.3 Spatial Distribution

#### Visualization

Recall that entries *u_p,v_* of *m* × *n* matrix *U* indicate the prevalence of clone *v* in biopsy *p* with spatial coordinates σ(*p*) = (*x, y*). Given the prevalence of a clone *v* in different biopsies and threshold *τ*, our goal is to draw isolines showing the clone at prevalence *τ* on a regular grid (A1). To that end, we first define an *X* × *Y* regular grid that can accommodate each biopsy location. We use bilinear interpolation to infer clonal prevalences for grid points that do not correspond to biopsies. As such, each point on the grid represents either a prevalence observed in a biopsy (colored black) or an interpolated value between biopsies (colored gray, see Fig. 3). Subsequently, we use the open source MarchingSquares.js^2^ D3-based implementation of the marching squares algorithm (Lorensen *et al*., 1987) to draw isolines.

**Fig. 3:**
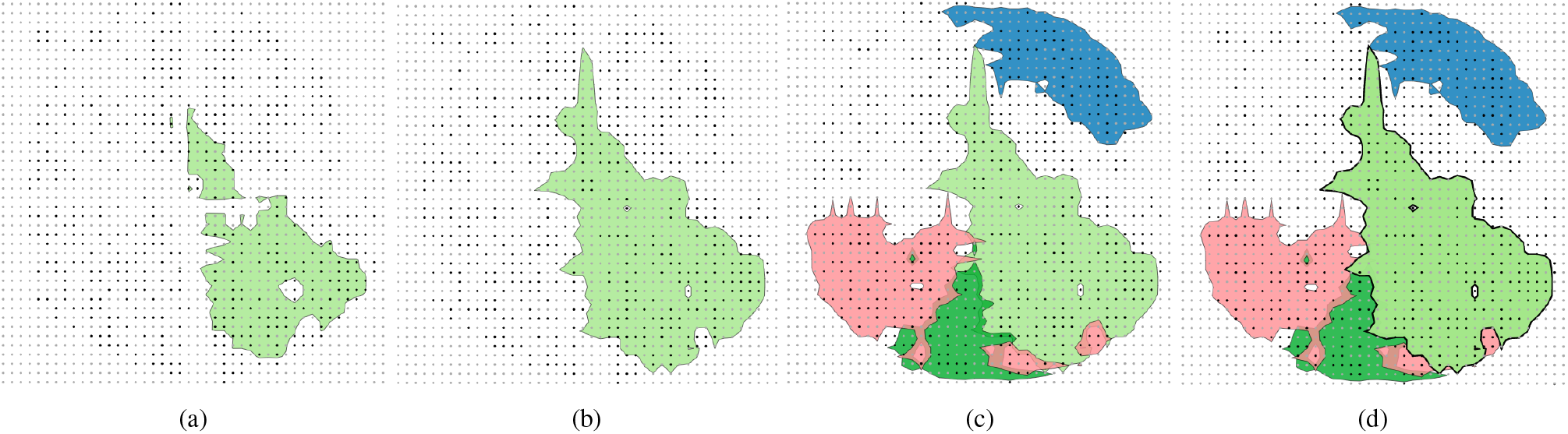
Visualization of spatial distribution of tumor clones. (a) We construct a regular grid accommodating the sequenced biopsies (black), followed by interpolation to obtain clonal prevalences at unsequenced grid points (gray). Given a threshold *τ* = 65%, we draw a set of isolines at which the clone occurs at the specified prevalence. Then, we fill the area enclosed by the set of isolines such that each grid point inside the filled area contains the clone at a prevalence of at least *τ*. (b) Decreasing the prevalence threshold *τ* to 20% results in a larger area. (c-d) Hover and click interactions bring corresponding boundary lines to the front of the grid.

For a matrix *U* with *n* clones, we draw *n* sets of clone-specific isolines, each set corresponding to a clone *v* and threshold *τ_v_*. The set of isolines corresponding to each clone is assigned a different color from a qualitative color palette from ColorBrewer^3^. To facilitate the spatial analysis of multiple clones, we fill the area enclosed by each set of clone-specific isolines using a transparent color. As such, the filled area is composed of grid points at which clone *v* occurs at a prevalence of *τ_v_* (Fig. 3a). Currently, up to 12 clones are supported with distinctive colors from ColorBrewer. More clones may be selected; however, the colors after the 12th clone may not be qualitatively distinctive.

#### Interactivity

The user is able to set the prevalence threshold via a slider for either individual clones (A5) or all clones simultaneously (A6) by selecting them in the phylogenetic tree. Toggling the threshold sliders will change the contour of the corresponding clonal boundaries on the grid (Fig. 3b). Hovering over a clonal boundary will highlight its corresponding vertex in the phylogeny tree, by thickening stroke-width on both elements, and vice versa (Fig. 3c-d). Additionally, this will bring the clone to the front of the grid. Clicking on a clone in the grid will “select” the clone, fix it to the foreground on the grid, bring up the corresponding clone-specific threshold slider, and increase the stroke width of the corresponding vertex in the phylogenetic tree. Hovering over a grid point will display a tool tip with the sample name at that coordinate.

### 3.4 Additional Tasks

In line with task A4, the user may restrict the analysis to a subset *Z* of biologically or clinically relevant mutations. Upon specifying *Z*, the phylogenetic tree and the grid will be redrawn using a new set of clones that are defined in terms of *Z*. Specifically, we intersect the mutations contained in each original clone with *Z*. Clones with the same set of resulting mutations will be merged and their clonal prevalences summed for each grid point (Fig. 2b). It is important to note that this feature allows the user to identify the presence of clones of interest on the visualization grid. In collapsing non-selected clones, each vertex on the phylogenetic tree will represent a group of clones, rather than an individual clone.

Following task A7, the user may export an SVG file containing the phylogenetic tree and/or the grid.

### 3.5 ClonArch

ClonArch is a web-based tool that implements the functionality described above using `D3`^4^. The input to ClonArch is a JSON file, containing both the set 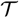 of phylogenetic trees and the frequency matrix *F*. ClonArch is open source and is available at: https://github.com/elkebir-group/ClonArch.

## 4 Results

### 4.1 Analysis of a Hepatocellular Carcinoma Tumor using ClonArch

We demonstrate how ClonArch enables one to study the spatial clonal architecture of tumors using a recent hepatocellular carcinoma (HCC) dataset (Ling *et al*., 2015). Ling *et al*. (2015) sequenced a total of 286 spatial biopsies in addition to a nearby matched normal sample. In their initial analysis, the authors identified 269 single-nucleotide variants (SNVs) in a subset of 23 whole-exome sequenced biopsies. They designated 209 SNVs as fixed, i.e. occurring in every tumor clone. From the remaining SNVs, Ling *et al*. (2015) selected a subset of 35 SNVs for targeted sequencing in the remaining 286−23 = 263 biopsies, yielding a variant allele frequency (VAF) for each of the 35 SNVs in each of the 263 biopsies. The authors then discretized the obtained VAFs and used a combination of neighbor joining and maximum parsimony to infer a phylogenetic tree *T*. Finally, the authors manually constructed a map of the clonal architecture of the tumor. Using the discretized frequencies, ClonArch is able to faithfully recreate the manually-constructed tumor spatial figure from Ling *et al*. (2015) on a grid (Fig. 4b), providing additional interactivity.

**Fig. 4:**
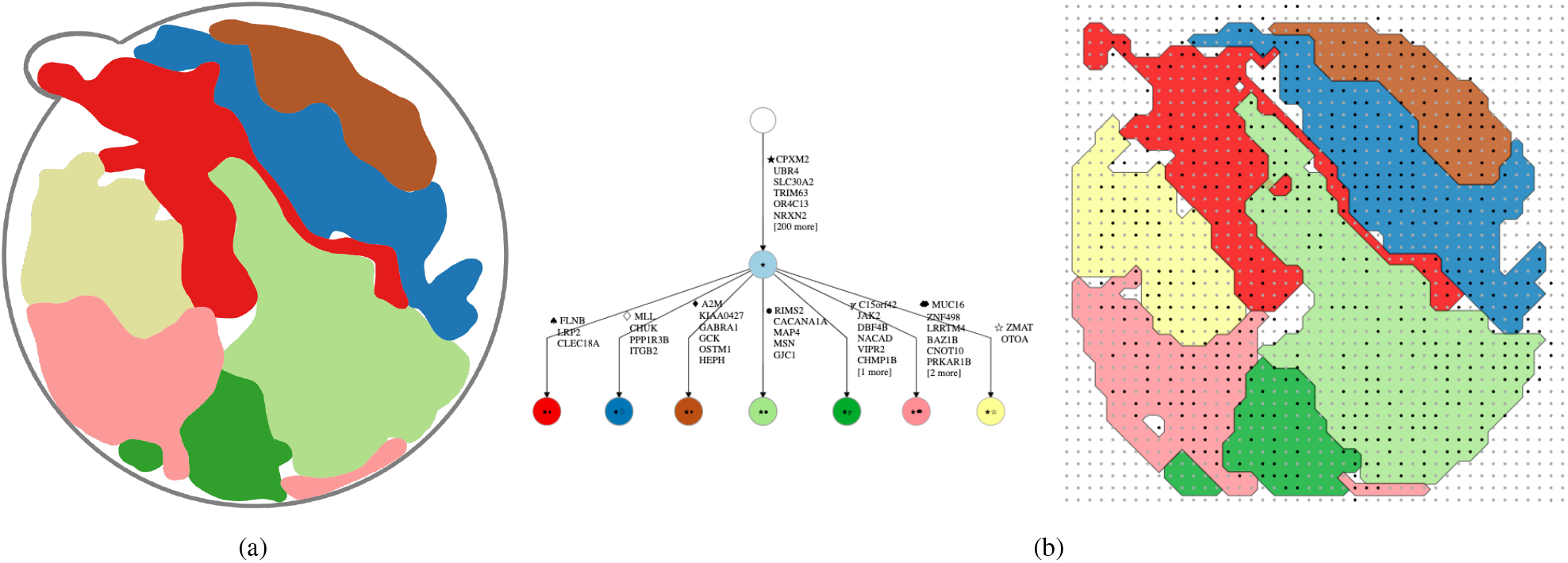
ClonArch automates the process of visualizing the spatial clonal architecture of a tumor. (a) Ling *et al*. (2015) manually visualized the spatial architecture of an HCC tumor using discretized mutation frequencies and a tree obtained by neighbor joining and maximum parsimony in disctinct biopsies of the tumor (figure adapted from Ling *et al*. (2015)).(b) ClonArch’s automated reconstruction of the figure with added interactivity, alongside the corresponding phylogenetic tree.

In the following, we show that ClonArch does not require one to discretize VAFs and clonal prevalences, enabling a more fine-grained and interactive view of the spatial clonal architecture.

#### Data preparation

Recall that ClonArch takes as input a set 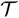 of phylogenetic trees and a frequency matrix *F*. For each tree 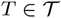, matrix *U* describes the clonal prevalence of each biopsy on a regular grid. While Ling *et al*. (2015) inferred T from discretized VAFs 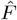, they did not infer *U*. In addition to describing how we computed *U* from *T* and *F*, we will describe how we obtained prevalences for each point on a regular grid.

Inspecting the provided tree *T* reveals that SNVs are homoplasy-free, i.e. are introduced exactly once on the tree and never subsequently lost. To simplify inference of *U*, we restrict our attention to copy-neutral SNVs on autosomes. SNVs present in copy-neutral autosomal regions have at most two chromosomal copies in each tumor cell. Under the assumption of no-homoplasy, at most one of these two copies will contain the SNV, amounting to a maximum VAF of 0.5. Inspection of *F* reveals that VAFs are ≤ 0.5 with the exception of 6 SNVS (OTOA, CLEC18A, GJC1, CHMP1B, PLEKHG6, OSTM1). In addition to excluding these 6 SNVs, we excluded 3 SNVs on the X chromosome (HEPH, ZMAT1, MSN), yielding frequency matrix *F*.

Next, we manually fit a regular grid to all sequenced biopsies, resulting in a 41 × 43 grid. After drawing a circular boundary with clonal prevalences set to 0, we used bilinear interpolation to infer missing values in *U*. This resulted in a total of about 400 real data points (a sequencing biospy may be covered by multiple grid points) and 1100 empty data points that were interpolated. Implementation details of both steps are in a Jupyter notebook on the git repository.

Assuming that the remaining 26 SNVs are copy-neutral and using an average of 0.7 purity obtained by multiplying the average VAFs of the trunk mutations by 2 (Ling *et al*., 2015) for the WES samples, we solved a linear program on the interpolated matrix to compute a frequency matrix *F* that best explains SNV frequencies. Running the tree enumeration tool SPRUCE (El-Kebir *et al*.,2016) on values from our interpolated frequency matrix *F* yields a set 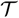 of two trees that describe our data and includes the tree reported by Ling *et al*. (2015).

#### Use case

In the following use case, we used ClonArch to analyze the spatial clonal architecture of this tumor, taking as input the set 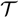 of trees and interpolated frequency matrix *F*. A screencast of this use case is availableat https://elkebir-group.github.io/ClonArch/screencast.html.

i. Phylogenetic tree overview. Initially, ClonArch shows a phylogenetic tree according to the default clones selected by the user (Fig. 5a-b). On hovering over a tree vertex, ClonArch shows a list of mutations at the corresponding clone (Fig. 2a). This enables the user to explore the phylogenetic relationships between clones (A2).
ii. Exploring the spatial distribution of different clones. At the same time, ClonArch shows a grid portraying the spatial distributions of tumor clones (Fig. 5a), in line with analysis task A1. We observe similarities between clones in the discretized reconstruction of Ling *et al*. (2015)‘s figure (Fig. 4a) and the spatial plot with real frequencies; namely, mutation clusters containing MUC16, MLL and CHUK, and RIMS2 appear in similar regions of the grid. Clones that are displayed on the grid correspond to the clones activated by the user in the phylogenetic tree. That is, clicking on different vertices in the tree toggles the visualization of the corresponding clones on the grid. To avoid clutter, we select only five clones to show on the grid; these are the vertices with incoming edges labeled with mutations: RIMS2, C15orf42 and JAK2, FLNB, GCK, and PPP1R3B. This version of the grid is much more straightforward; each clone is clearly distinguished on the grid (Fig. 5b-c).
iii. Hovering over grid points. To further explore the spatial distributions of these six clones, we take a closer look at the grid points they are drawn on. The different color of grid points indicates whether the data point was from a biopsy (black), or interpolated (gray). Hovering over the grid points also brings up a tool tip that specifies whether the point is from a biopsy, and if so, the name of the sample (Fig. 5d). This information helps gauge the validity of a boundary shape. An oddly-shaped clonal boundary is less likely to be an artifact of interpolation if it contains a fairly distributed amount of real biopsy samples. Following our example, the shape of the dark green clone (C15orf42 and JAK2) seems to be well-supported by an even distribution of real samples (Fig. 5c). In contrast, the upper boundary region of the light green clone (RIMS2) has irregular spikes towards the top and left sides that contain interpolated values; it seems likely that this is an artifact of interpolation (Fig. 5c).
iv. Selecting driver mutations. Next, we select a subset of interesting mutations to look at (A4). Ling *et al*. (2015) reported six driver mutations in this dataset: CPXM2, TMPRSS13, DNAH7, TSC1, TP53, and CCAR1,. Selecting these mutations updates the phylogenetic tree, revealing that all are fixed mutations (Fig. 5e). That is, they are present in every clone in the sample. In line with analysis task A5, we may adjust the prevalence threshold to observe the distribution of the clones containing these driver mutations at different levels (Fig. 5e).
v. Selecting additional mutations. To add more clones onto the grid, we can select additional mutations to closely examine. In general, the selection of mutations will be driven by biological and clinical reasons, such as the driver status of the mutation or its neo-antigenic potential. Here, besides the six drivers, CCAR1, CPXM2, DNAH7, TMPRSS13, TP53, and TSC1, we add two more mutation clusters: MLL and CHUK, and PPP1R3B (Fig. 5f). With multiple clones now on the grid, we may choose to examine either clone-specific or global prevalence at different thresholds (A6).
vi. Exploring different phylogeny to describe spatial data. In cases where mutation frequencies are ambiguous, we may find that multiple phylogenetic trees describe the data. ClonArch can visualize the spatial architecture of tumors according to different phylogenetic trees, allowing us to potentially disambiguate between multiple trees (A3). Fig. 5f visualizes PPP1R3B with its parent, MLL and CHUK. The child clone is completely overlapped by the parent, which illustrates a likely scenario. Meanwhile, Fig. 5g represents a second tree that also describes this data; here, PPP1R3B is a child of the root. Distribution of clones on the grid show that the child clone does not overlap with the parent at all. Spatial analysis suggests the tree illustrated in Fig. 5f is a more likely hypothesis.
vii. Saving results. Finally, we download the resulting grid and phylogenetic tree as SVG files (A7). These elements may be useful later as a reference, or a point of comparison to a new dataset or different subset of clones/mutations.

**Fig. 5:**
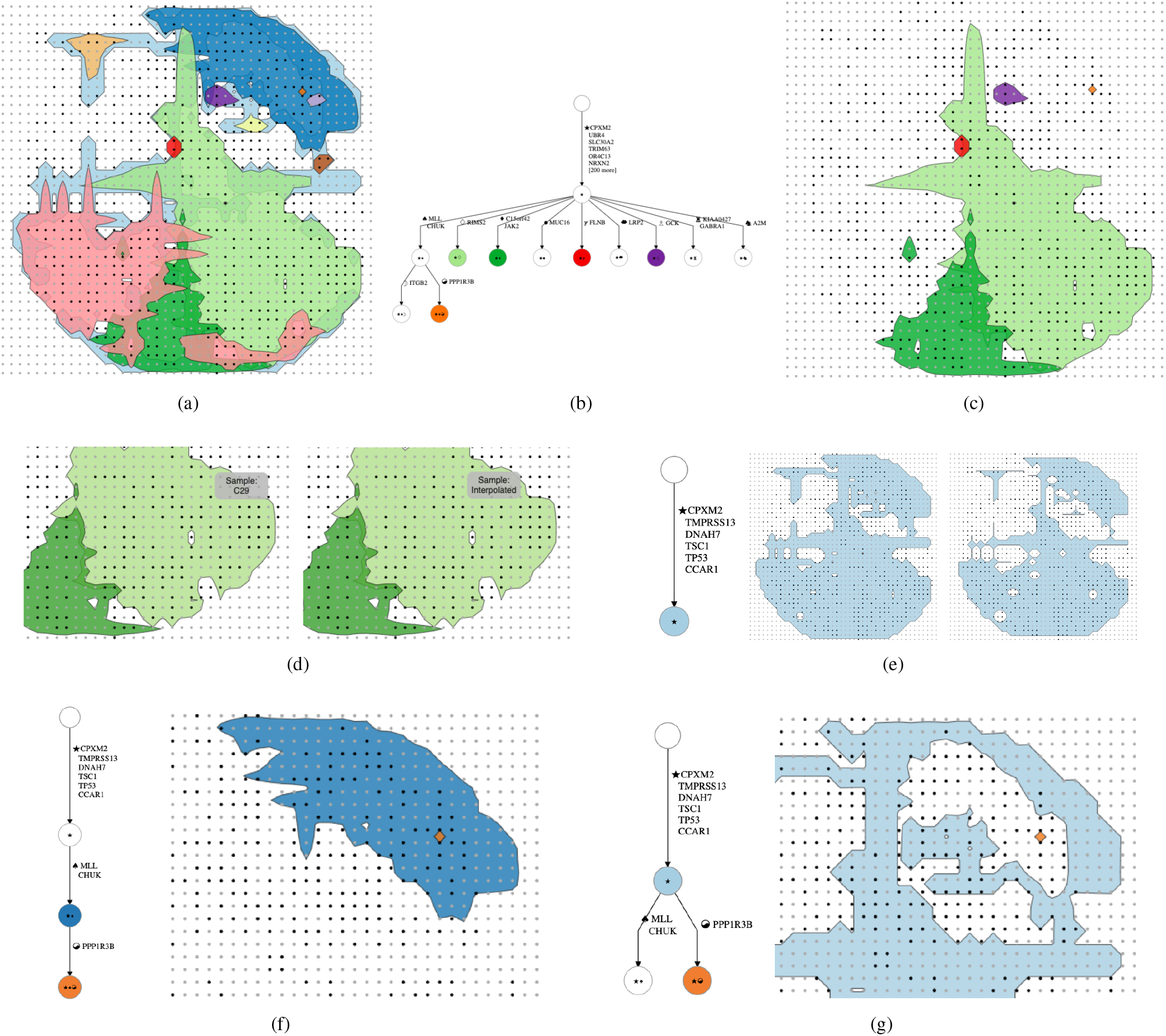
A case study using ClonArch to analyze a hepatocellular carcinoma dataset. (a) Initially, all vertices on the selected tree are visualized on the grid. (b-c) Clicking on vertices on the tree will hide their corresponding clone to make the grid less cluttered. (d) Hovering over grid points will reveal the origin of the data point (either the sample name or interpolated). (e) The phylogenetic tree reveals that all driver mutations in this study are fixed—they occur in every clone; we can adjust their prevalence threshold to see their presence throughout the tumor. (f) Here, we observe our clone of interest, PPP1R3B, and its parent, MLL and CHUK, from the original tree. We can see that, spatially, the child (PPP1R3B) is closely related to its parent. (g) The same clone, PPP1R3B, is visualized on a different tree, with its parent as the root. In this example, PPP1R3B does not occur very closely to its parent. Therefore, we believe that the tree illustrated in (f) is a more likely hypothesis

### 4.2 Simulations

Although ClonArch successfully visualizes the spatial clonal architecture assuming a large number of biopsies, we asked the question: can this tool be applicable to smaller datasets? Applying a simulation that mimics the invasive glandular model of tumor growth (Noble *et al*., 2019), we generated trees and frequency matrices at grid sizes of 3 × 3 and 5 × 5 (Fig. 6a-b). We observe that bilinear interpolation in the marching squares algorithm will lead to more inaccurate delineations of clonal compositions with few biopsies; this is apparent in our 3 × 3 example (Fig. 6a).

**Fig. 6:**
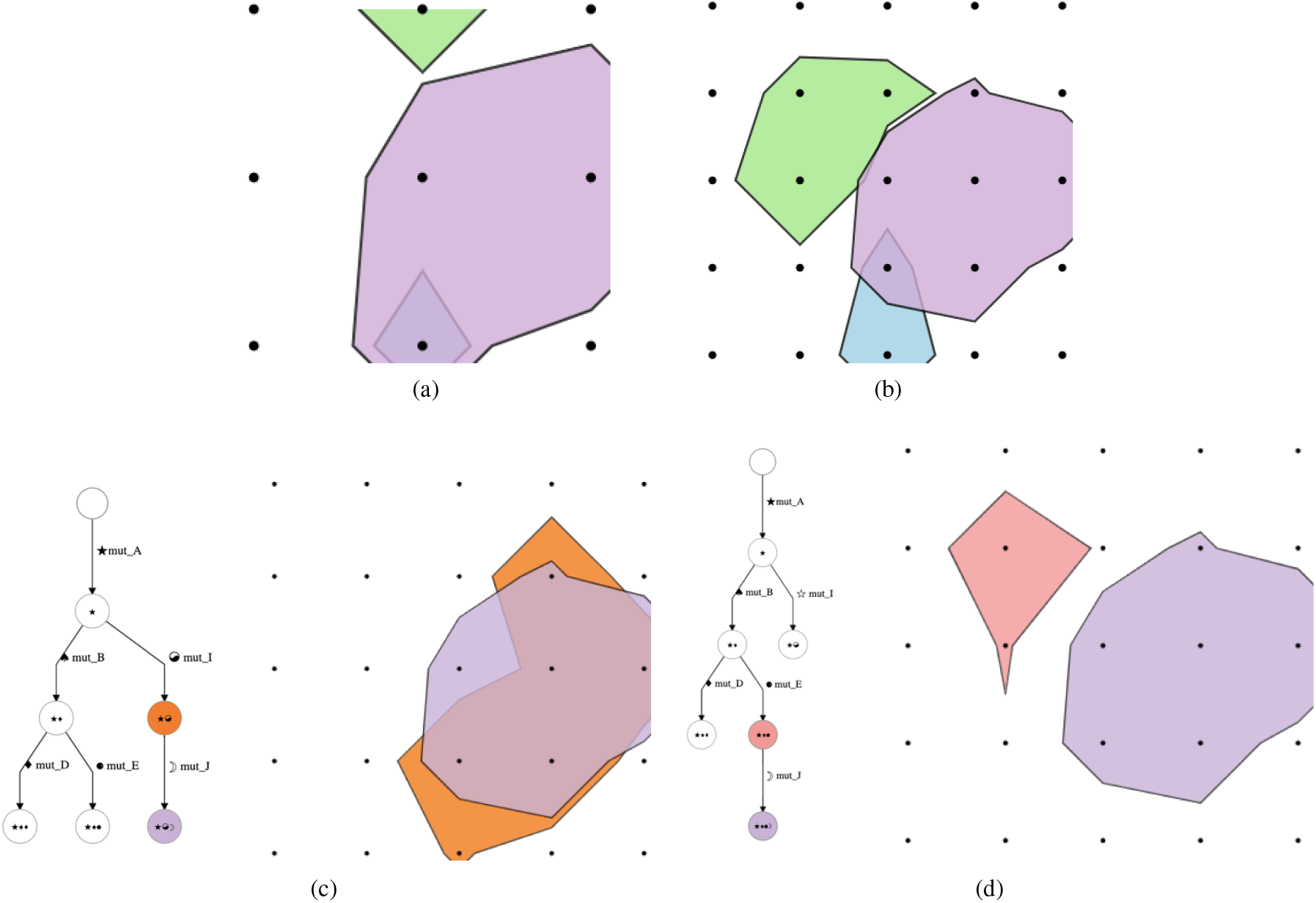
Simulations show that applicability of ClonArch increases with increasing number of biopsies. (a) A simulated 3 × 3 grid. We see that bilinear interpolation in the marching squares algorithm leads to more inaccurate delineations of clonal compositions with a fewer biopsies. (b) A simulated 5 × 5 grid shows that spatial structure can be identified with less than 25 samples. (c) With the 5 × 5 simulated example, we attempt to use ClonArch to disambiguate between two trees. Here, we see mut_J and its parent, mut_I, according to its tree. The two clones are overlapping with eachother, showing that it was very likely for mut_J to have originated from mut_I. (d) Here, we see mut_J and its parent, mut_E, as according to the tree. Unlike the last example, however, the two clones do not overlap, making this phylogenetic tree a less likely hypothesis.

We also observe that with less than 25 samples, as shown in the 5 × 5 grid, we begin to see the spatial structure of the tumor. Fig. 6c-d shows a simulated example with ambiguity between two trees. In the first tree (c), we see that mut_J is a child of mut_I; in the visualized grid, mut_J and mut_I share a high amount of overlap, indicating that mut_J could have originated from mut_I. In the second tree (Fig. 6d), mut_J is a child of mut_E. However, the two clones do not share any overlap in their spatial architecture, making it more difficult to believe that mut_J could have originated from mut_I. Therefore, the first tree (Fig. 6c) seems to be the more likely hypothesis in this scenario. ClonArch helps us disambiguate between multiple trees by allowing users to analyze tumor spatial patterns according to different phylogeny.

## 5 Discussion

In this work, we addressed the lack of available visualization tools for studying the spatial aspects of intra-tumor heterogeneity. We introduced ClonArch, a method to interactively visualize the spatial distribution of clones in a tumor given a set of phylogenetic trees and mutation frequencies at distinct biopsies. Accounting for multiple trees elucidates the consequences of non-uniqueness of solutions on spatial clonal composition and distribution. In particular, ClonArch facilitates the prioritization of trees in terms of consistency between evolutionary relationships of clones and their spatial locations.

We used the marching squares algorithm to draw enclosed boundaries around clones above a specified threshold at each spatial location. The analysis tasks outlined in the requirements section guided our design and implementation of ClonArch. These decisions were motivated by the hepatocellular carcinoma case study (Ling *et al*., 2015). Although the HCC dataset contains a large number of biopsies, simulated data illustrates that ClonArch can successfully visualize spatial structure of smaller datasets as well.

Despite these contributions, ClonArch has some limitations. First, the current interface works best with about 12 clones before the visualization starts to look cluttered, and colors become less distinct. This is an inherent limitation of the chosen visualization approach, and a tradeoff for accommodating for more clones. To resolve the limitation of clutter in the future, a zooming and panning feature may be implemented to view regions in both the tree and spatial grid close up. Second, data preparation can be complicated. Shaping the spatial data into a form that can be computationally analyzed is not straightforward. Currently, empty grid points have to be interpolated, and a boundary of zeros around the tumor has to be specified for proper interpolation. In future work, we will address this limitation by developing user-friendly scripts that automate the process of fitting/interpolating a regular grid given spatial coordinates of sequencing biopsies. In particular, we plan to develop convex-hull based algorithms to avoid interpolation artefacts.

Third, ClonArch requires many biopsies to provide an accurate spatial representation. As we observed with simulated data, we ideally require a 5 × 5 grid. Presently, few available datasets have a sufficient number of samples, thus only one dataset covering one cancer type (HCC) has been analyzed. However, we observe that recent studies are beginning to sequence increasing numbers of tumor biopsies, especially in the context of understanding spatial heterogeneity (Ding *et al*., 2019; Alves *et al*., 2019), which ClonArch will be able to visualize. We expect the number of such high-resolution spatial datasets to increase with decreasing costs in sequencing technology. Additionally, though our application treats biopsies locations as 2D coordinates, tumors are actually 3D in nature. The depth of the biopsy needle needs to be accounted for in order to generate a full, accurate model of the tumor. Ding *et al*. (2019) have utilized a 3D sampling approach; each tumor is evenly sliced, and then spatially organized samples are collected from each slice.

To facilitate 3D visualizations, one future direction might be to extend ClonArch using the marching cubes algorithm. Fourth, our method currently only visualizes a single tumor mass. Leveraging recent work in migration analysis of metastatic cancers (El-Kebir *et al*., 2018), it will be of interest to extend our tool to visualize clonal patterns of metastatic spread given biopsies from matched primary and metastasis samples. Finally, while incorporating information on phylogeny and spatial location, our method loses the temporal aspect that other cancer visualizations include. Implementing this feature in the future requires biopsies from a tumor mass at varying slices and differing time points; then, we can explore the tumor using a slider to view the time axis. In the future, spatial visualizations may be required to support 4 dimensions—3D space and time—to truly represent the spatiotemporal clonal dynamics of tumors as close as possible.

## Footnotes

1 https://github.com/dagrejs/dagre-d3

2 https://github.com/RaumZeit/MarchingSquares.js

3 http://colorbrewer2.org

4 https://d3js.org

## References

Alves, J. M. et al. (2019). Rapid evolution and biogeographic spread in a colorectal cancer. Nature Communications, 10(1), 5139.

Deshwar, A. G. et al. (2015). PhyloWGS: Reconstructing subclonal composition and evolution from whole-genome sequencing of tumors. Genome Biology, 16(1), 35.

Ding, X. et al. (2019). Genomic and epigenomic features of primary and recurrent hepatocellular carcinomas. Gastroenterology, 157(6), 1630–1645.e6.

El-Kebir, M. (2018). SPhyR: tumor phylogeny estimation from single-cell sequencing data under loss and error. Bioinformatics, 34(17), i671–i679.

El-Kebir, M. et al. (2015). Reconstruction of clonal trees and tumor composition from multi-sample sequencing data. Bioinformatics, 31(12), i62–i70.

El-Kebir, M. et al. (2016). Inferring the Mutational History of a Tumor Using Multi-state Perfect Phylogeny Mixtures. Cell Systems.

El-Kebir, M. et al. (2018). Inferring parsimonious migration histories for metastatic cancers. Nature Genetics, 50(5), 718–726.

Fisher, R. et al. (2013). Cancer heterogeneity: implications for targeted therapeutics. British journal of cancer, 108(3), 479–485.

Gerlinger, M. et al. (2012). Intratumor heterogeneity and branched evolution revealed by multiregion sequencing. N Engl J Med, 366(10), 883–92.

Gerlinger, M. et al. (2014). Genomic architecture and evolution of clear cell renal cell carcinomas defined by multiregion sequencing. Nat Genet, 46(3), 225–33.

Jahn, K. et al. (2016). Tree inference for single-cell data. Genome biology, 17(1), 86.

Jamal-Hanjani, M. et al. (2017). Tracking the Evolution of Non-Small-Cell Lung Cancer. New England Journal of Medicine, 376(22), 2109–2121.

Krzywinski, M. (2016). Visualizing Clonal Evolution in Cancer. Molecular Cell.

Kvitek, D. J. and Sherlock, G. (2013). Whole Genome, Whole Population Sequencing Reveals That Loss of Signaling Networks Is the Major Adaptive Strategy in a Constant Environment. PLOS Genetics.

Ling, S. et al. (2015). Extremely high genetic diversity in a single tumor points to prevalence of non-darwinian cell evolution. PNAS.

Lorensen, W. E. et al. (1987). Marching cubes: A high resolution 3D surface construction algorithm, volume 21. ACM.

Malikic, S. et al. (2015). Clonality inference in multiple tumor samples using phylogeny. Bioinformatics, 31(9), 1349–1356.

Mamlouk, S. et al. (2017). Dna copy number changes define spatial patterns of heterogeneity in colorectal cancer. Nature Communications, 8(1), 14093.

Miller, C. A. et al. (2014). Sciclone: Inferring clonal architecture and tracking the spatial and temporal patterns of tumor evolution.

Miller, C. A. et al. (2016). Visualizing tumor evolution with the fishplot package for r. BMC Genomics.

Noble, R. et al. (2019). Spatial structure governs the mode of tumour evolution. bioRxiv.

Nowell, P. C. (1976). The clonal evolution of tumor cell populations. Science, 194(4260), 23–8.

Popic, V. et al. (2015). Fast and scalable inference of multi-sample cancer lineages. Genome biology, 16(1), 91.

Pradhan, D. and El-Kebir, M. (2018). On the Non-uniqueness of Solutions to the Perfect Phylogeny Mixture Problem. RECOMB.

Ross, E. M. and Markowetz, F. (2016). OncoNEM: inferring tumor evolution from single-cell sequencing data. Genome biology, 17(1), 69.

Roth, A. et al. (2014). PyClone: statistical inference of clonal population structure in cancer. Nature methods, 11(4), 396–398.

Roth, A. et al. (2016). Clonal genotype and population structure inference from single-cell tumor sequencing. Nature methods, 13(7), 573–576.

Schroeder, M. P. et al. (2013). Visualizing multidimensional cancer genomics data.

Smith, M. A. et al. (2017). E-scape: interactive visualization of single-cell phylogenetics and cancer evolution. Nature Methods.

Tabassum, D. P. and Polyak, K. (2015). Tumorigenesis: it takes a village. Nature Reviews Cancer, 15(8), 473–483.

